# Improved two-photon imaging of GPCR-based optogenetic neurotransmitter sensors using orthogonally polarized excitation

**DOI:** 10.1101/2021.11.15.468606

**Authors:** Mauro Pulin, Kilian E. Stockhausen, Olivia Andrea Masseck, Martin Kubitschke, Björn Busse, J. Simon Wiegert, Thomas G. Oertner

## Abstract

Fluorescent proteins such as GFP are best excited by light that is polarized parallel to the dipole axis of the fluorophore. In most cases, fluorescent proteins are randomly oriented, resulting in unbiased images even when polarized light is used for excitation, e.g. in two-photon microcopy. Here we reveal a surprisingly strong polarization sensitivity in a class of GPCR-based neurotransmitter sensors where the fluorophore is anchored on both ends. In tubular structures such as dendrites, this effect led to a complete loss of membrane signal in dendrites running parallel to the polarization direction of the excitation beam. Our data reveal a major problem for two-photon measurements of neurotransmitter concentration that has not been recognized by the neuroscience community. To remedy the sensitivity to dendritic orientation, we designed an optical device that generates interleaved pulse trains of orthogonal polarization, removing the orientation bias from images. The passive device, which we inserted in the beam path of an existing two-photon microscope, also removed the strong direction bias in second harmonic generation (SHG) images. We conclude that for optical measurements of transmitter concentration with GPCR-based sensors, orthogonally polarized excitation is essential.

## 1. Introduction

Excitation of fluorescence depends on the orientation of the fluorophore with respect to the polarization of the excitation light. The emitted photons are also polarized, an effect known as fluorescence anisotropy. This effect can be exploited to measure the speed of rotation of fluorescent proteins in solution [1,2], for example to detect interactions during molecular signaling. Commercial two-photon microscopes use linearly polarized light for excitation. In the case of two-photon light-sheet microscopy, where fluorescence is detected orthogonally to the excitation sheet, it has been noted that better results are achieved when the direction of polarization is optimized [3]. In classical point-scanning 2p microscopes, however, the direction of polarized excitation is ignored, as fluorophores in biological preparations are typically freely diffusing and thus randomly oriented.

To detect the release of neurotransmitters and neuromodulators in intact brain tissue, numerous genetically encoded sensors have been developed. A very successful strategy is to splice circularly permuted GFP (cpGFP) into G protein-coupled neurotransmitter receptors (GPCRs), which resulted in several highly specific sensors with a large dynamic range [4–8]. GPCRs have seven transmembrane domains, and the cpGFP is inserted into the third intracellular loop, tethering both sides of the GFP barrel to transmembrane helices of the receptor. Upon ligand binding, movements of the helices are transferred to the cpGFP, changing its fluorescence. The relative change in fluorescence (df/f_0_) depends on neurotransmitter concentration in a logistic fashion. However, depending on the length and rigidity of the linker sections, tethering cpGFP to transmembrane domains may orient the barrel parallel to the membrane.

Here we show that two-photon imaging of GPCR-based sensors is surprisingly sensitive to the orientation of the imaged dendrite relative to the polarization of the excitation beam. As a consequence, entire sections of reporter neurons can be non-fluorescent, which makes it impossible to draw conclusions about the spatial distribution of neurotransmitter release from the relative change in fluorescence. To solve this problem, we developed a passive optical device that produces two collinear output beams with orthogonal polarization (‘X-Pol’). When inserted into the beam path of a two-photon microscope, the X-Pol device removes the orientation bias when imaging membrane proteins. As the polarization sensitivity of genetically encoded sensors cannot be known *a priori*, we suggest using orthogonal polarized pulse trains as a new standard for two-photon microscopy.

## 2. Materials and Methods

### Organotypic slice cultures

Hippocampal slice cultures from Wistar rats of either sex were prepared at postnatal day 4–6. Rats were anesthetized with 80% CO_2_ 20% O_2_ and decapitated. Hippocampi were dissected in cold slice culture dissection medium containing (in mM): 248 sucrose, 26 NaHCO_3_, 10 glucose, 4 KCl, 5 MgCl_2_, 1 CaCl_2_, 2 kynurenic acid, 0.001% phenol red (saturated with 95% O_2_, 5% CO_2_, pH 7.4). Tissue was cut into 400 μM thick sections on a tissue chopper (McIlwain) and cultured on membranes (Millipore PICMORG50) at 37° C in 5% CO_2_. The slice culture medium contained (for 500 ml): 394 ml Minimal Essential Medium, 100 ml heat inactivated donor horse serum, 1 mM L-glutamine, 0.01 mg ml^−1^ insulin, 1.45 ml 5M NaCl, 2 mM MgSO_4_, 1.44 mM CaCl_2_, 0.00125% ascorbic acid, 13 mM D-glucose. Wistar rats were housed and bred at the University Medical Center Hamburg-Eppendorf. All procedures were performed in compliance with German law and according to the guidelines of Directive 2010/63/EU. Protocols were approved by the Behörde für Gesundheit und Verbraucherschutz of the City of Hamburg.

### Single cell electroporation

At DIV 13-17, CA1 neurons in rat organotypic hippocampal slice culture were transfected by single-cell electroporation [9]. Thin-walled borosilicate pipettes (~10 MΩ) were filled with plasmid DNA encoding the GFP-based sensors along with a cytoplasmic red fluorescent protein tdimer2 diluted in intracellular solution to 50 ng μl^−1^ and 10 ng μl^−1^, respectively. Pipettes were positioned against neurons under visual control (IR-DIC) and DNA was ejected using an Axoporator 800A (Molecular Devices) with 50 hyperpolarizing pulses (−12 V, 0.5 ms) at 50 Hz. Imaging experiments were conducted 2-4 days after electroporation at room temperature (21-23°) in artificial cerebrospinal fluid (ACSF) containing (in mM) 135 NaCl, 2.5 KCl, 2 CaCl_2_, 1 MgCl_2_, 10 Na-HEPES, 12.5 D-glucose, 1.25 NaH_2_PO_4_ (pH 7.4).

### Darken sensor

We developed an improved serotonin sensor named Darken (darkening 5-HT_1A_ receptor-based sensor). The sensor design is based on dLight_1_ [4]. Due to its high sensitivity, the native human 5-HT_1A_ receptors was used as the sensing scaffold. We replaced the third intracellular loop of 5-HT_1A_ with a circular mutated form of GFP (cpGFP) from GCaMP6 [10] flanked by mutated linker sequences. Linker length is identical to dLight_1.2_. Darken is bright in the unbound form and decreases its fluorescence upon serotonin binding.

### Bone sections

Healthy femoral bone was obtained during autopsy at the Department of Forensic Medicine at the University Medical Center Hamburg-Eppendorf (Hamburg, Germany) from a 44-year old organ donor in line with previously published protocols (PV 3486) [10]. Femoral cross sections from the mid-diaphysis were cut using a diamond saw (EXAKT Advanced Technologies GmbH) and fixed in 3.7% formaldehyde for 3 days. The sample was dehydrated in an increasing alcohol series and embedded undecalcified in glycolmethacrylate (Technovit 7200, Heraeus Kulzer GmbH). Using an automatic grinding machine (EXACT Advances Technologies GmbH), the sample was ground to a thickness of 100 μm.

### Two-photon microscopy

The custom-built two-photon imaging setup was based on an Olympus BX51WI microscope with LUMPlan W-IR2 60× 1.0 NA objective. A galvanometric scanner (6215H, Cambridge Technology) was controlled by the open-source software package ScanImage [11]. Two pulsed Ti:Sapph lasers (MaiTai DeepSee, Spectra Physics) tuned to 930 nm and controlled by electro-optic modulators (350-80, Conoptics) were coupled at orthogonal polarization to excite GFP and tdimer2. In some experiments, we activated one or the other to generate images at orthogonal polarization (Fig. 2). For all other experiments, the polarization of a single Ti:Sapph laser was either rotated by 90° using a half-wave plate (690–1080 nm achromatic; Thorlabs) or split into orthogonally polarized pulse trains using the X-Pol device. Emitted photons were collected through the objective and oil-immersion condenser (1.4 NA, Olympus) with two pairs of photomultiplier tubes (H7422P-40, Hamamatsu) [12]. 560 DXCR dichroic mirrors and 525/50 and 607/70 emission filters (Chroma) were used to separate green and red fluorescence. Excitation light was blocked by short-pass filters (ET700SP-2P, Chroma).

### Image analysis

Two-photon image stacks were de-interleaved (red/green channel) and z-projected (maximum intensity projection, MIP). Displayed images are MIPs; magenta and green levels were adjusted separately and linearly (Gamma = 1). Intensity profiles along the x-axis of a rectangular region of interest placed across the apical dendrite were generated with Fiji/ImageJ 1.53c and plotted with GraphPad Prism 8.4.3.

### X-Pol device

The prototype device used in this study is based on the 30 mm cage system (Linos, Thorlabs) and can be rotated around the optical axis. It consists of the following parts:

| ITEM | DESCRIPTION | Cat No. | QTY |
| --- | --- | --- | --- |
| 1 | Polarizing Beamsplitter Cube, 10 mm, N-BK7, 980 nm | Edmund Optics #48-880 | 2 |
| 2 | Right-Angle Prism Mirror, Protected Silver, L = 10 mm | Thorlabs MRA10-P01 | 2 |
| 3 | Counter Support (mounting platform for optics) | Linos G061161000 | 2 |
| 4 | Solid Cage Plate (drill and tap to accept mirror mount) | Linos G061008000 | 1 |
| 5 | Compact Kinematic Mirror Mount | Thorlabs KMSS/M | 1 |
| 6 | 30 mm Cage Plate, SM1-Threaded | Thorlabs CP08/M | 2 |
| 7 | Cage Assembly Rod, 2" long, Ø 6 mm | Thorlabs ER2 | 4 |
| 8 | SM1 Lens Tube | Thorlabs SM1L10 | 2 |
| 9 | Clamp for SM1 Lens Tubes | Thorlabs SM1TC | 2 |
| 10 | Ø12.7 mm Optical Post, M6 Tap, L = 50 mm | Thorlabs TR50/M | 2 |
| 11 | Ø12.7 mm Pedestal Post Holder, L = 54.7 mm | Thorlabs PH50E/M | 2 |
| 12 | Clamping Fork, 31.5 mm counterbored slot | Thorlabs CF125C/M | 2 |
| 13 | Ring-Actuated Iris Diaphragm (for alignment) | Thorlabs SM1D12D | 1 |

### Alignment procedure

The device was inserted with a 45° tilt before the beam expander and centered on the excitation laser beam using the entrance pinhole (iris diaphragm) and an infrared scope. Axial orientation is not very critical as the device basically acts as a parallel plate. Some angular deviation of the exit beam was noted due to the limited parallelism of the beam splitting cubes (+/− 3 arcmin). This deviation was compensated by adjusting the following beam steering mirrors (periscope). The mirror prism inside the device was coarsely adjusted so that a single output spot was produced on an infrared viewer card held at a distance of about 1 m. Imaging a fluorescent structure in focusing mode, the mirror prism was fine adjusted to achieve maximum image brightness, corresponding to perfect overlap of the PSFs generated by the orthogonally polarized beams. This is best achieved when the person adjusting the mirror prism is able to see the live image on the computer screen. Lastly, we adjusted the pulse pre-compensation (MaiTai DeepSee) to account for the added GVD of 20 mm N-BK7 glass (1400 fs^2^). Conveniently, rotating the device in its holders to horizontal (0°) or vertical (90°) orientation switches off its beam splitting function, resulting in output polarization (and repetition rate) that is identical to the input beam. In working position (near 45°), some power (~12%) is lost at the second, beam-combining cube. This is due to imperfect polarization of the reflected beam in the first, beam-splitting cube. For safety, the exit of this light must be blocked, either with a black metal plate or by a Si-photodiode to measure laser power. Due to this power loss in the reflected sub-beam, the ideal 50/50 splitting position is ~55° rotation, not 45°. The position can be optimized by temporarily blocking the reflected sub-beam inside the device, which should drop the output power to exactly 50%.

## 3. Results

We transfected hippocampal CA1 neurons in organotypic slice culture [13] by single-cell electroporation [9] to express the following membrane-bound cpGFP-based sensors together with the red cytoplasmic label tdimer2 [14]: The GPCR-based dopamine sensor dLight_1.2_[4], the norepinephrine sensor GRAB_NE1h_ [7], a newly developed serotonin sensor (Darken) based on dLight_1_, and the glutamate sensor iGluSnFR [15]. For comparison, we also expressed GPI-anchored GFP [16] together with tdimer2 in some cells. Coupling two identical Ti:Sapph lasers tuned to 930 nm with a polarizing beam splitter cube allowed us to image the same structure using two different (orthogonal) polarization directions. For better comparison, we selected cells with vertically oriented apical dendrites for imaging.

The dopamine sensor dLight_1.2_ contains a circularly permuted GFP inserted between transmembrane domain 5 and 6 of the D1 dopamine receptor, orienting the GFP barrel parallel to the plasma membrane. For optical measurements, we added 10 μM dopamine to the extracellular solution to maximize the brightness of dLight_1.2_. Fluorescence of dLight_1.2_ was strongly dependent on the direction of laser polarization (Fig. 1a-c). Dendrites orthogonal to the direction of laser polarization were fluorescent while parallel ones were not. In contrast, polarization direction had no effect on the red fluorescence signal of (freely diffusible) tdimer2. When we compared dLight_1.2_ fluorescence intensity in response to dopamine application, we observed a strong dopamine-dependent fluorescence increase for dendrite-normal polarization (background-subtracted df/f_0_ = 123%), but no signal for dendrite-parallel polarization (Fig. 1d). Thus, using linear polarized excitation, it is not possible to relate the relative changes in dLight_1.2_ fluorescence to dopamine concentration changes.

**Fig. 1.**
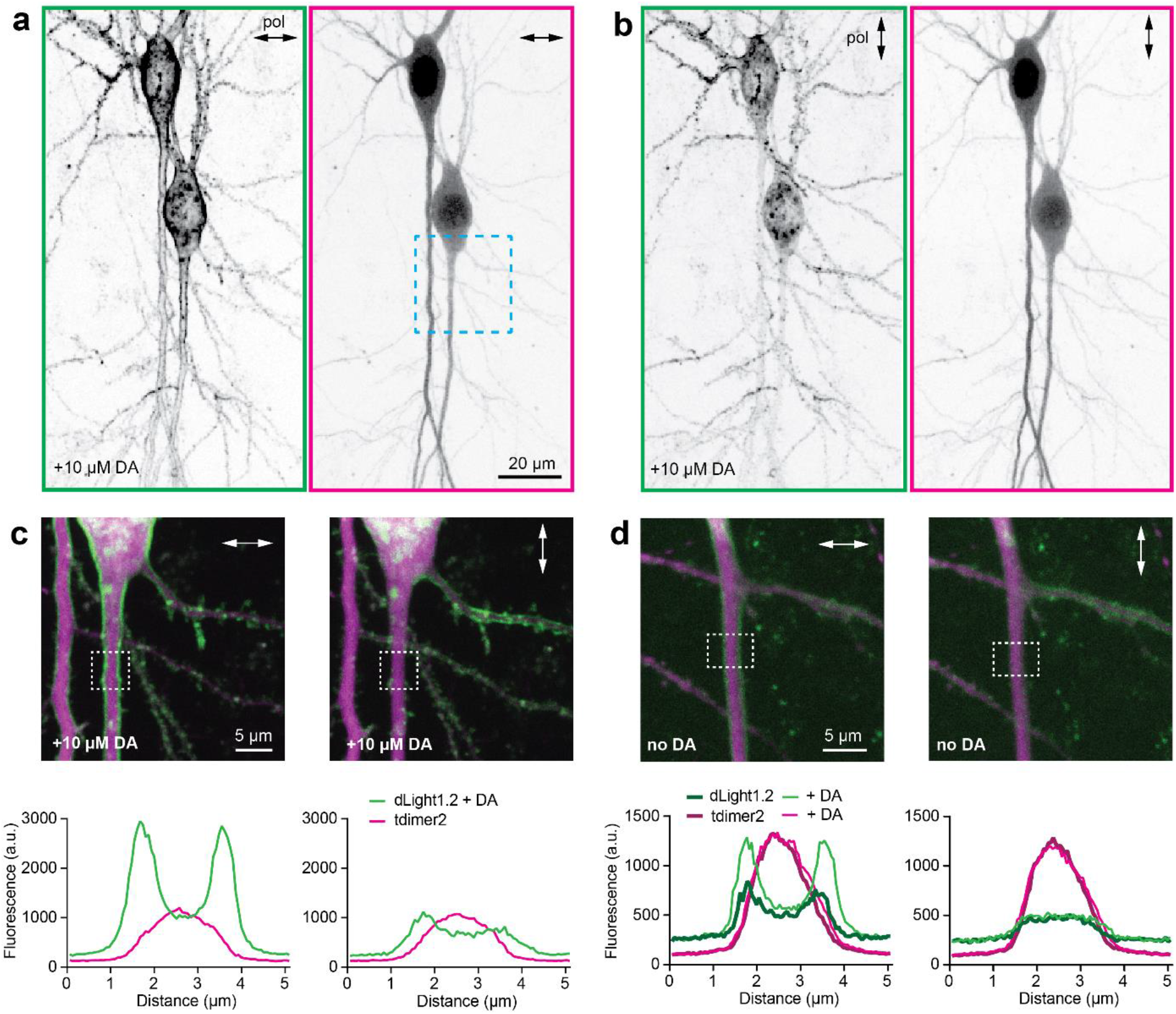
Effect of laser polarization (930 nm) on the brightness of GPCR-based dopamine sensor dLight_1.2_. (a) Laser polarization (arrow) orthogonal to the apical dendrites of CA1 pyramidal cells excites bright fluorescence of dLight_1.2_ (left) and cytoplasmic tdimer2 (right). Images are contrast-inverted maximum intensity projections. dLight_1.2_ was saturated with dopamine (10 μM). Apical dendrite (blue box) is enlarged in panel c. (b) Laser polarization (arrow) parallel to the apical dendrites excites no fluorescence of dLight_1.2_ in the apical dendrites (left) while cytoplasmic fluorescence (right) is unchanged. (c) Fluorescence profiles across the vertically oriented apical dendrite (same neuron as a, b; 10 μM dopamine). Laser polarization orthogonal to dendrite excites membrane-localized green fluorescence of dLight_1.2_ (left). Polarization parallel to dendrite excites little green fluorescence while red (cytoplasmic) fluorescence is unchanged (right). Note that dLight_1.2_ fluorescence in horizontally oriented branches shows the opposite polarization-dependence. (d) Application of 10 μM dopamine strongly increases dLight_1.2_ fluorescence when laser polarization is normal to dendrite (left). Dopamine application had no effect on fluorescence when laser polarization was parallel to the dendrite (right).

Next, we tested Darken, a genetically encoded sensor for serotonin we recently developed. It is based on the architecture of dLight_1.2_, but decreases its fluorescence in response to serotonin. Thus, we could visualize bright images of Darken-expressing CA1 pyramidal cells in the absence of the agonist serotonin (Fig. 2a). Again, we observed no membrane-localized green fluorescence when the polarization of the laser was parallel to the dendritic membrane (Fig. 2b). Diffuse green fluorescence in the cytoplasm likely results from sensor molecules that were not trafficked to the surface as well as immature (green fluorescent) tdimer2.

**Fig. 2.**
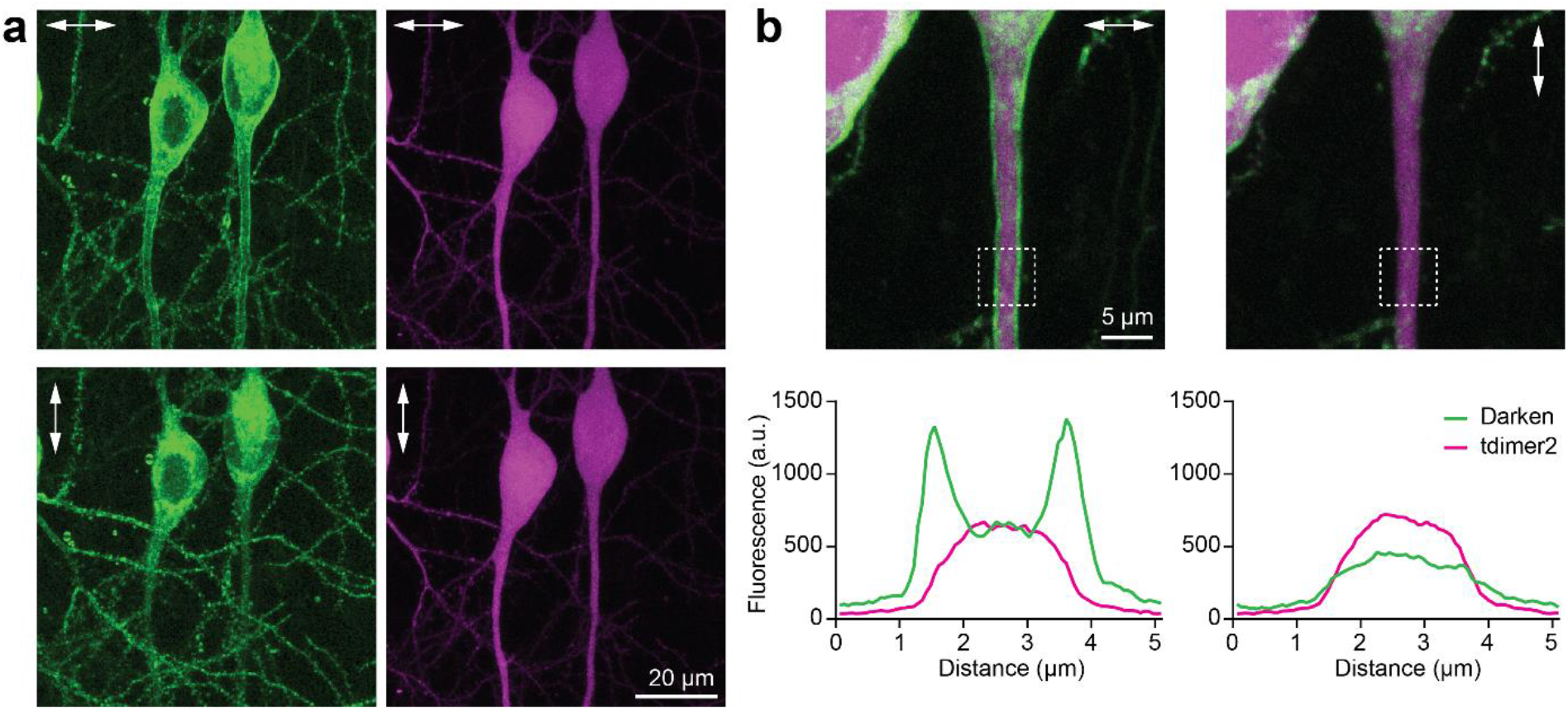
Effect of laser polarization (930 nm) on the brightness of GPCR-based serotonin sensor Darken. (a) Laser polarization normal to the apical dendrites of CA1 pyramidal cells excites bright fluorescence of Darken (left) and cytoplasmic tdimer2 (right). Laser polarization parallel to the apical dendrites excites no fluorescence of Darken in the apical dendrites (left) while cytoplasmic fluorescence (right) is unchanged. (b) Fluorescence profiles across the apical dendrite (same neuron as a). Polarization normal to dendrite shows membrane-localized green fluorescence of Darken (left), polarization parallel to dendrite shows no membrane-localized fluorescence.

To change the polarization of excitation, we used two different methods: We either coupled two separate lasers tuned to the same wavelength at orthogonal polarization, activating one or the other (Fig. 2), or we rotated the polarization of a single laser by 90° using a half-wave plate (Fig. 1, Fig. 3). The results were identical, confirming that the direction of polarization determined which parts of the neuronal membrane were visible.

**Fig. 3.**
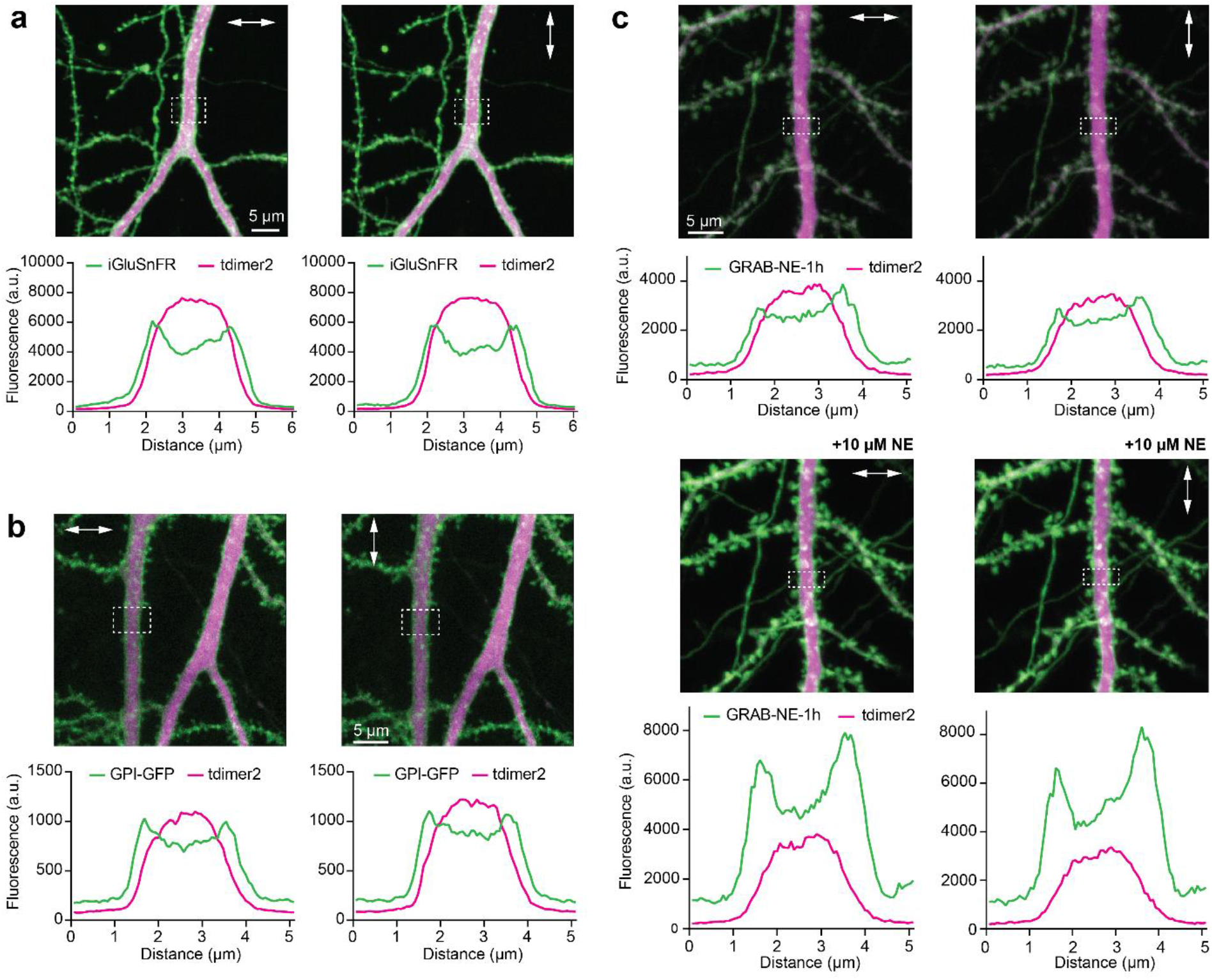
Examples of membrane-anchored GFP constructs that are not polarization-sensitive. Dotted boxes mark analyzed regions. (a) The glutamate sensor iGluSnFR was not sensitive to polarization direction. (b) Membrane-anchored GFP (GPI-GFP) was not sensitive to polarization direction. (c) Norepinephrine sensor GRAB_NE1h_ did not show strong polarization dependence of baseline fluorescence (without agonist). Addition of norepinephrine (NE) led to a strong increase in fluorescence independent of polarization direction.

Interestingly, not all membrane-tethered GFP constructs are sensitive to laser polarization. The glutamate sensor iGluSnFR, which is anchored to the membrane via a single transmembrane segment, did not change its fluorescence intensity when excited at different polarization directions (Fig. 3a), and neither did GFP tethered to the membrane via glycosylphosphatidylinositol (GPI-GFP, Fig. 3b). We conclude that single-sided anchors leave sufficient degrees of freedom to randomize the angle of the fluorophore with respect to the membrane plane. GRAB_NE1h_ is a GPCR-based, high affinity norepinephrine sensor with cpGFP attached to transmembrane segments on both ends, but in contrast to dLight_1.2_ and Darken, it retains a large part of the third intracellular loop as a flexible linker. When expressed in CA1 pyramidal cells, GRAB_NE1h_ did show a strong fluorescence increase in response to noradrenaline application, but no polarization dependence (Fig. 3c).

For quantitative measurements of neurotransmitter concentrations *in vivo*, the orientation-sensitivity of some of the GPCR-based sensors is a problem. Coupling two lasers tuned to the same wavelength at orthogonal polarization is possible, but not very economical. We designed a compact device that splits the beam of a single laser into two sub-beams of equal power (50/50) but orthogonal polarization, and recombines the sub-beams collinearly (Fig. 4a-c). The device uses two polarizing beam-splitter cubes with 980 nm antireflection coating. To minimize group velocity dispersion (GVD), we used cubes made of n-BK7, rather than the denser N-SF1 (Flint) glass. For the same reason, we used silver coated prisms to direct the reflected beam through air instead of using total internal reflection in right angle prisms. The device is mounted in circular clamps so it can be rotated around the beam to 45° (for 50/50 splitting) or used horizontally (no beam splitting). At 45° (the ‘on’ position), it works for horizontally and vertically polarized input beams, creating a stream of output of pulses with alternating polarization (45°, 225°). Both horizontal (0°) and vertical (90°) orientation of the device result in an unchanged output beam (‘off’ positions).

**Fig. 4.**
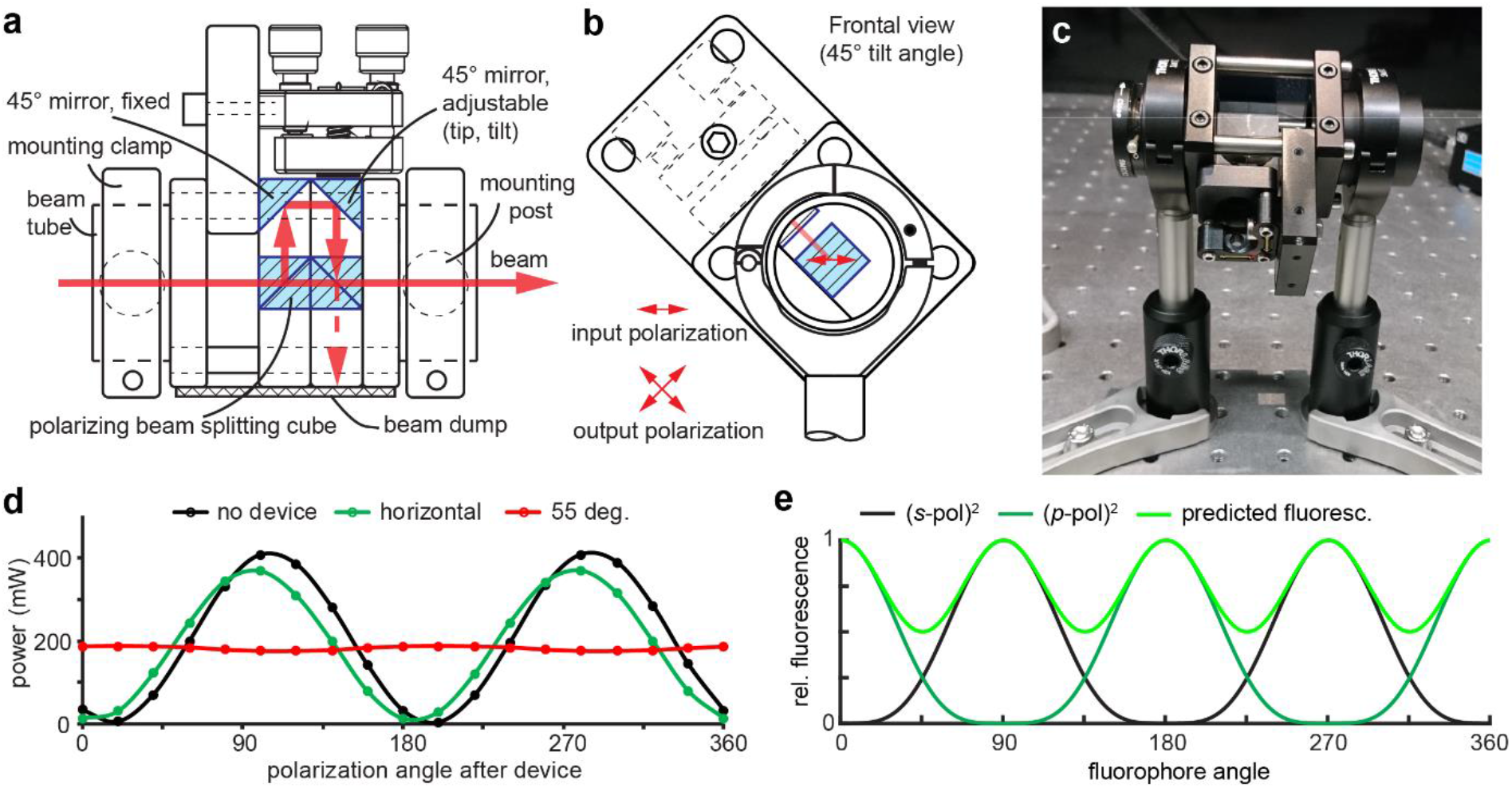
X-Pol device to produce orthogonal pulse trains. (a) Schematic drawing of the beam path through the X-pol device. One of the mirror prisms is adjustable to align the output beams. (b) In operation, the device is tilted 45° with respect to the polarization direction of the input beam. (c) Prototype mounted at 45° tilt. (d) Laser power at different polarization directions. Inserting the device leads to a small power loss due to surface reflections (green). The total output power is independent of polarization angle at a tilt angle of ~55° (red curve), due to some power loss of the reflected beam at the 2^nd^ (recombining) cube. (e) Simulation of fluorescence-dependence on fluorophore orientation (with X-Pol device). As two-photon excitation is proportional to the square of the pulse power, and total fluorescence is the sum of emission from molecules excited by *s*-polarized and *p*-polarized pulses, respectively, we predict a residual orientation-dependence of fluorescence.

We inserted a beam splitting cube in a rotating mount to analyze the average power after the X-pol device at different polarization directions (Fig. 4d). With the X-Pol device in the ‘off’ position (horizontal), the output beam was still horizontally polarized. When the device was tilted around the optical axis, it started splitting the beam into orthogonal components, reaching a flat output spectrum at a tilt angle of ~55°. In this position, a power loss of ~12% occurred at the device (at 930 nm), mostly due to the mixed polarization of the reflected beam exiting the first beam splitter cube. The horizontally polarized component is not reflected at the second beam splitting cube; it crosses the cube and is lost. Therefore, slightly more power has to be directed into the reflected path to achieve a balanced output, which in our prototype was reached at 55° tilt angle. For safety reasons, lateral exit of the ‘lost’ beam has to be prevented. In our prototype, it is absorbed by a strip of matte black aluminum. With a side-mounted Si photodiode, it could be utilized to measure the power of the imaging beam.

Although we achieved a flat distribution of laser power across all polarization directions, two-photon excitation of (perfectly aligned) fluorophores would still be expected to show some polarization dependence (Fig. 4e). Two-photon excitation depends on the excitation power squared, and molecules oriented at 45° to the two orthogonally polarized pulse trains are excited at twice the frequency, but with half the power per pulse. Note that pulses cannot interact; they have a temporal offset of ~70 ps (20 mm in air). In our biological preparations, we did not see any direction sensitivity with the X-Pol device inserted, most likely because the alignment of GFP dipoles (or in the case of SHG imaging, collagen fibrils) is not perfect.

We tested the X-Pol device first on the bright serotonin sensor Darken, which was completely invisible on dendrites aligned with the polarization direction of the laser, but perfectly visible on membranes orthogonal to the polarization direction (Fig. 5a). Inserting the device in the beam path at 55° tilt angle achieved the same results than combining two separate lasers with orthogonal polarization: Green fluorescence was bright along the entire membrane, independent of dendritic orientation (Fig. 5b). The less efficient excitation when pulses with half the energy are used at twice the frequency was apparently compensated by the larger fraction of fluorophores available for excitation. As a result, using the same average laser power, the intensity of red fluorescence of cytoplasmic tdimer2 is near identical in images 5a and 5b. Note that the green/red ratio is higher in thinner dendrites as it reflects the surface-to-volume ratio of the compartment.

**Fig. 5.**
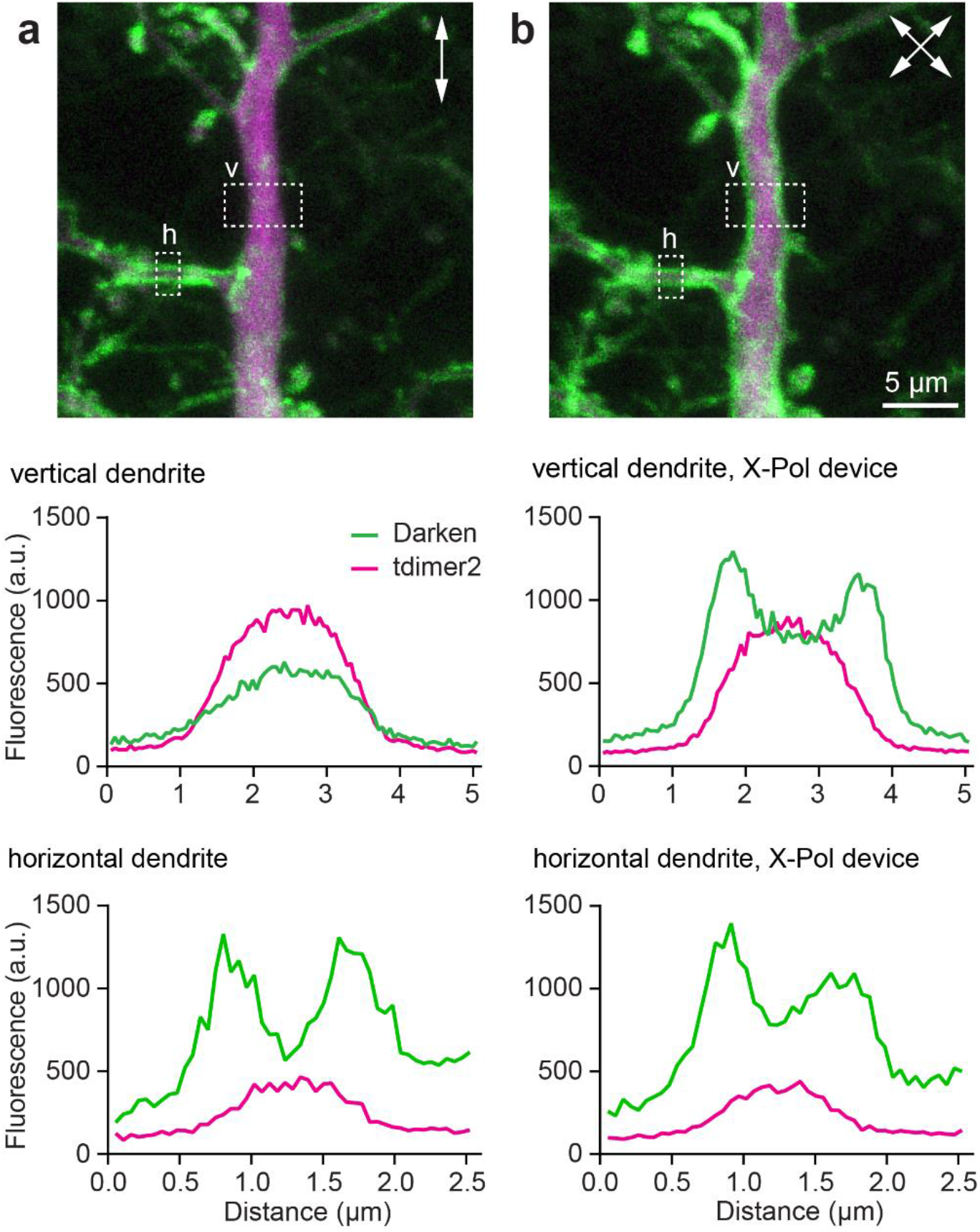
CA1 pyramidal cell apical dendrite expressing the serotonin sensor Darken (green) and cytoplasmic tdimer2 (red). (a) X-Pol device oriented horizontally (no beam spitting) rendered Darken on the apical dendrite (vertical, v) invisible, while there was bright Darken fluorescence on horizontal dendrites (h). (b) X-Pol device at 55° resulted in excellent visibility of Darken on all dendrites. Note that the red (cytoplasmic) fluorescence is weaker in thinner dendrites due to their small volume.

We tested the X-Pol device also in label-free imaging of thin sections from human bone. Oriented collagen fibers generate a strong second harmonic signal when oriented parallel to the polarization direction of the excitation laser, revealing details of fiber organization below the diffraction limit of the microscope [17]. We used polarization sensitive second harmonic generation (P-SHG) to investigate the lamellar organization of osteons (Fig. 6a). In standard configuration, the two-photon microscope did not generate green emission from the sectors of the osteon where its lamellae were orthogonal to the polarization, resulting in two dark sectors. The tilt of the dark sectors in our images reflects a 15° tilt of the galvanometric x-mirror’s axis of rotation (XY Galvo Block, Cambridge Technology), resulting in a 75° polarization angle in the images.

**Fig. 6.**
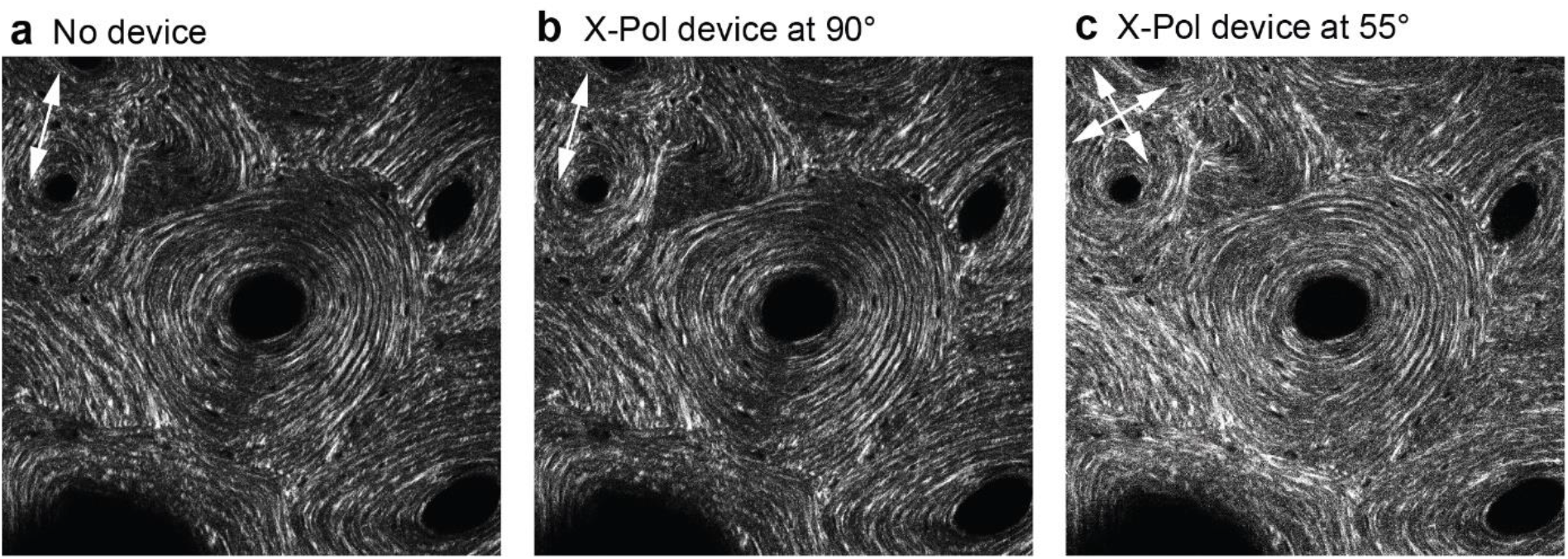
Second harmonic generation in a bone section at 1020 nm excitation. (a) Image of an osteon shows dark sectors where collagen fibers are oriented orthogonal to the polarization direction of the IR laser. (b) Insertion of X-Pol device into the beam path in vertical direction (90°) has little effect on the second harmonic image. (c) Turning the device to 55° results in orthogonally polarized sub-beams. Note the absence of dark sectors.

Inserting the X-Pol device into the beam path (Fig. 6b) and rotating it to 55° with respect to the laser polarization (Fig. 6c) resulted in evenly distributed signals from all sectors of the osteon, removing the artifact produced by linear polarization. To compensate for light loss inside the polarization device, laser power was increased by ~20% (Fig. 6c).

Osteons provide a convenient test sample as their lamellar collagen organization is known to be rotationally symmetric. The strong polarization dependence of SHG images is even more problematic in tissues that have not the same degree of crystalline organization (e.g. skin and tissue), as the absence of signal in such images can be mistakenly interpreted as absence of collagen. Mamillary tumors, for example, are characterized by an increased collagen density, and two-photon SHG imaging has been successfully used to visualize such malignancies [18,19]. As we show, orthogonal polarization excitation makes SHG images easier to interpret, which could be helpful for the investigation and diagnosis of cancer.

## 4. Discussion

We report unexpectedly strong effects of polarized excitation in two-photon microscopy of a particular type of GFP-labeled membrane proteins, namely GPCR-based neurotransmitter sensors. No polarization dependence was observed for the soluble red fluorescent protein tdimer2. Similarly, flexible, single-ended tethering, as in GPI-anchored GFP and iGluSnFR, did not result in measurable anisotropy. If the fluorophore can rotate in a cone, its dipole orientation relative to the direction of polarization is randomized. To obtain polarization-sensitive probes with single-ended tether, the linker has to form a continuous α-helix to prevent angular movements [20].

In GPCR-based neurotransmitter sensors, the GFP barrel is inserted into the third intracellular loop and thus tethered at both ends (Fig. 7). This architecture orients the GFP barrel more or less parallel to the cell membrane, depending on the length of the linkers [21]. Based on available structural information [20,22–24], we propose that in highly anisotropic sensors, the dipole moment of cpGFP is oriented normal to the membrane plane (Fig. 7). Interestingly, we could not detect significant polarization dependence of the norepinephrine sensor GRAB_NE1h_. It has been pointed out that in these kind of GRAB sensors, large parts of the third intracellular loop are retained and provide a flexible link to the GFP barrel [21]. Thus, subtle differences in sensor design can have profound effects on the sensitivity to polarization. In the development of sensors intended for *in vivo* imaging, typically performed with two-photon microscopes using polarized excitation, isotropic fluorescence should indeed be a selection criterion. In confocal microscopes, lasers are usually fiber-coupled and thus lose their polarization before reaching the sample, which is ideal for imaging fluorescent proteins.

**Fig. 7.**
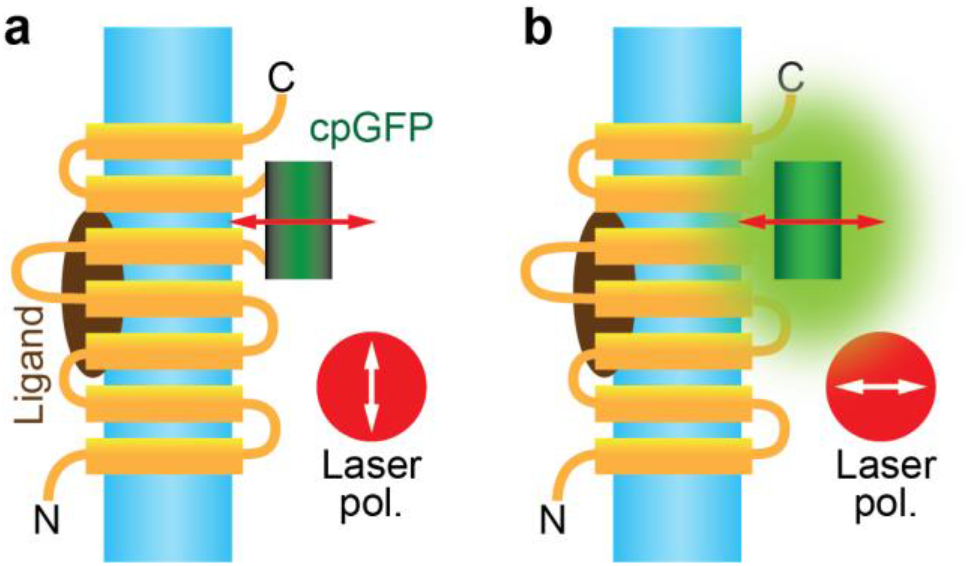
Proposed dipole orientation in polarization-sensitive GPCR sensors. (a) Laser polarization parallel to the dendritic membrane (blue) does not excite cpGFP. (b) Laser polarization must be aligned with the dipole of cpGFP (red arrow) to excite fluorescence. Note that rotation of the molecule in the membrane plane does not affect dipole orientation.

For quantitative neurotransmitter measurements deep in in brain tissue, we propose a simple solution, to alternate pulses of orthogonal polarization. We show that the X-Pol device significantly reduces direction bias in neurons. For specific applications, fluorescence-detected linear dichroism could also be exploited, e.g. to visualize physical interactions between subunits of G proteins [25]. However, this works best in spherical cells and requires an active device to rapidly switch laser polarization. For the class of GPCR-based neurotransmitter sensors, the goal is to precisely measure extracellular neurotransmitter concentrations on the entire surface of the neuron. As we demonstrate, this goal is put in jeopardy if linear polarized excitation is used.

Our prototype X-Pol device was assembled from off-the-shelf components and is therefore inexpensive to reproduce. We would like to point out, however, that precise alignment of the optical components inside the device is important to ensure complete overlap of the two point-spread functions generated by the sub-beams. If the mirror prisms are not well adjusted, loss of resolution or even double images may occur. Aligning the device to the center of the beam is less critical as long as beam clipping is avoided. Our prototype has a generous free aperture (10 mm) and can thus be inserted before or after a 3x beam expander. The added dispersion of 20 mm N-BK7 glass (*GVD* = 70 fs^2^/mm) was readily compensated by pre-chirping (MaiTai DeepSee, SpectraPhysics). Flint glass (N-SF1), which is typically used for beamsplitting cubes, induces more GVD (247 fs^2^/mm). To further reduce GVD, broadband thin plate beamsplitters could be arranged at Brewster’s angle in a chevron design, again mounted with axial tilt to achieve 50/50 splitting [26]. Instead of rotating the X-Pol device, the polarization of the incoming beam could be tilted with a half-wave plate. However, adding an achromatic half-wave plate roughly doubles the cost of the device without obvious benefits.

The X-Pol device doubles the pulse repetition rate while cutting the energy of individual pulses in half. This effect should require an increase in average power (by 2^0.5^) to achieve the same image intensity. On the other hand, more fluorophores can be excited when using orthogonal pulse trains, increasing the effective concentration of fluorescent proteins. We found that these two effects are in balance, and no increase in average power was required when imaging with the X-Pol device at 55°.

Increased repetition rates have been reported to reduce bleaching in biological preparations [27]. Some of the beneficial effects shown in that study, which used a more complex pulse splitter design, may have been due to orthogonally polarized output pulses and in consequence, recruitment of additional fluorophores.

Excitation with orthogonal polarization could also be advantageous for two-photon excitation of channelrhodopsin as well as opsin-derived sensors of membrane voltage [28]. Indeed, polarization-dependence of genetically encoded voltage sensors has recently been reported [29]. In membranes, cytoskeletal labeling and in fixed tissue, where the rotation of fluorophores is slowed down or locked, conventional linearly polarized two-photon microscopes continuously excite the same subset of fluorophores, leading to anisotropic photobleaching [30]. Orthogonally polarized pulse trains double the number of excitable fluorophores, increasing brightness (compared to excitation pulses of the same energy and repetition frequency polarized in one direction) or reducing bleaching when imaging with adapted laser power. Based on our experience, we predict that two-photon microscopy will follow the lead of confocal microscopy and use mixed polarization as default imaging mode.

## Funding

This study was supported by the German Research Foundation DFG through SPP 1665 (T.G.O), SPP 1926 (J.S.W.), SFB 936 (T.G.O. and J.S.W.) and FOR 2419 (T.G.O. and J.S.W.), MA 4692/6-2 (O.A.M.), and by ERC grants to J.S.W. (Starting grant) and to T.G.O. (Synergy grant).

## Acknowledgements

The authors would like to thank Iris Ohmert, Jan Schröder and Kathrin Sauter for excellent technical assistance. pAAV-hSyn-dLight1.2 was a gift from Lin Tian (Addgene plasmid # 111068), pAAV-hSyn-GRAB_DA1h (Addgene plasmid # 113050) and pAAV-hSyn-GRAB_NE1h (Addgene plasmid # 123309) were gifts from Yulong Li, iGluSnFR was a gift from Loren Looger (Addgene plasmid # 41732), GPI-GFP was a gift from Martin Heine, tdimer2 was a gift from Roger Y. Tsien.

## Disclosures

Portions of the technologies presented in the manuscript are patented or patent pending. However, the authors have no commercial interest, arrangement, or affiliation that could be perceived as a conflict of interest in the context of this manuscript.

## Data availability

Data underlying the results presented in this paper are not publicly available at this time but may be obtained from the authors upon reasonable request.

## Notes

### Competing Interest Statement

The authors have declared no competing interest.

## References

1. S. T. Hess, E. D. Sheets, A. Wagenknecht-Wiesner, and A. A. Heikal, “Quantitative analysis of the fluorescence properties of intrinsically fluorescent proteins in living cells,” Biophys. J. 85(4), 2566–2580 (2003).

2. A. H. A. Clayton, Q. S. Hanley, D. J. Arndt-Jovin, V. Subramaniam, and T. M. Jovin, “Dynamic fluorescence anisotropy imaging microscopy in the frequency domain (rFLIM),” Biophys. J. 83(3), 1631–1649 (2002).

3. G. de Vito, P. Ricci, L. Turrini, V. Gavryusev, C. Müllenbroich, N. Tiso, F. Vanzi, L. Silvestri, and F. S. Pavone, “Effects of excitation light polarization on fluorescence emission in two-photon light-sheet microscopy,” Biomed. Opt. Express 11(8), 4651 (2020).

4. T. Patriarchi, J. R. Cho, K. Merten, M. W. Howe, A. Marley, W.-H. Xiong, R. W. Folk, G. J. Broussard, R. Liang, M. J. Jang, H. Zhong, D. Dombeck, M. von Zastrow, A. Nimmerjahn, V. Gradinaru, J. T. Williams, and L. Tian, “Ultrafast neuronal imaging of dopamine dynamics with designed genetically encoded sensors,” Science 360(6396), (2018).

5. F. Sun, J. Zeng, M. Jing, J. Zhou, J. Feng, S. F. Owen, Y. Luo, F. Li, H. Wang, T. Yamaguchi, Z. Yong, Y. Gao, W. Peng, L. Wang, S. Zhang, J. Du, D. Lin, M. Xu, A. C. Kreitzer, G. Cui, and Y. Li, “A Genetically Encoded Fluorescent Sensor Enables Rapid and Specific Detection of Dopamine in Flies, Fish, and Mice,” Cell 174(2), 481–496.e19 (2018).

6. M. Jing, Y. Li, J. Zeng, P. Huang, M. Skirzewski, O. Kljakic, W. Peng, T. Qian, K. Tan, J. Zou, S. Trinh, R. Wu, S. Zhang, S. Pan, S. A. Hires, M. Xu, H. Li, L. M. Saksida, V. F. Prado, T. J. Bussey, M. A. M. Prado, L. Chen, H. Cheng, and Y. Li, “An optimized acetylcholine sensor for monitoring in vivo cholinergic activity,” Nat. Methods 17(11), 1139–1146 (2020).

7. J. Feng, C. Zhang, J. E. Lischinsky, M. Jing, J. Zhou, H. Wang, Y. Zhang, A. Dong, Z. Wu, H. Wu, W. Chen, P. Zhang, J. Zou, S. A. Hires, J. J. Zhu, G. Cui, D. Lin, J. Du, and Y. Li, “A Genetically Encoded Fluorescent Sensor for Rapid and Specific In Vivo Detection of Norepinephrine,” Neuron 102(4), 745–761.e8 (2019).

8. J. Wan, W. Peng, X. Li, T. Qian, K. Song, J. Zeng, F. Deng, S. Hao, J. Feng, P. Zhang, Y. Zhang, J. Zou, S. Pan, M. Shin, B. J. Venton, J. J. Zhu, M. Jing, M. Xu, and Y. Li, “A genetically encoded sensor for measuring serotonin dynamics,” Nat. Neurosci. 24(5), 746–752 (2021).

9. J. S. Wiegert, C. E. Gee, and T. G. Oertner, “Single-cell electroporation of neurons,” Cold Spring Harb. Protoc. 2017(2), 135–138 (2017).

10. P. Milovanovic, E. A. Zimmermann, C. Riedel, A. vom Scheidt, L. Herzog, M. Krause, D. Djonic, M. Djuric, K. Püschel, M. Amling, R. O. Ritchie, and B. Busse, “Multi-level characterization of human femoral cortices and their underlying osteocyte network reveal trends in quality of young, aged, osteoporotic and antiresorptive-treated bone,” Biomaterials 45, 46–55 (2015).

11. T. A. Pologruto, B. L. Sabatini, and K. Svoboda, “ScanImage: Flexible software for operating laser scanning microscopes,” Biomed Eng Online 2(1), 13 (2003).

12. T. G. Oertner, “Functional imaging of single synapses in brain slices,” Exp Physiol 87(6), 733–736 (2002).

13. C. E. Gee, I. Ohmert, J. S. Wiegert, and T. G. Oertner, “Preparation of slice cultures from rodent hippocampus,” Cold Spring Harb. Protoc. 2017(2), 126–130 (2017).

14. R. E. Campbell, O. Tour, A. E. Palmer, P. A. Steinbach, G. S. Baird, D. A. Zacharias, and R. Y. Tsien, “A monomeric red fluorescent protein,” Proc Natl Acad Sci U S A 99(12), 7877–7882 (2002).

15. J. S. Marvin, B. Scholl, D. E. Wilson, K. Podgorski, A. Kazemipour, J. A. Müller, S. Schoch, F. J. U. Quiroz, N. Rebola, H. Bao, J. P. Little, A. N. Tkachuk, E. Cai, A. W. Hantman, S. S.-H. Wang, V. J. DePiero, B. G. Borghuis, E. R. Chapman, D. Dietrich, D. A. DiGregorio, D. Fitzpatrick, and L. L. Looger, “Stability, affinity, and chromatic variants of the glutamate sensor iGluSnFR,” Nat. Methods 15(11), 936–939 (2018).

16. G. Kondoh, X. H. Gao, Y. Nakano, H. Koike, S. Yamada, M. Okabe, and J. Takeda, “Tissue-inherent fate of GPI revealed by GPI-anchored GFP transgenesis,” FEBS Lett. 458(3), 299–303 (1999).

17. J. C. Mansfield, V. Mandalia, A. Toms, C. Peter Winlove, and S. Brasselet, “Collagen reorganization in cartilage under strain probed by polarization sensitive second harmonic generation microscopy,” J. R. Soc. Interface 16(150), (2019).

18. T. Meyer, N. Bergner, C. Bielecki, C. Krafft, D. Akimov, B. F. M. Romeike, R. Reichart, R. Kalff, B. Dietzek, and J. Popp, “Nonlinear microscopy, infrared, and Raman microspectroscopy for brain tumor analysis,” J. Biomed. Opt. 16(2), 021113 (2011).

19. P. P. Provenzano, D. R. Inman, K. W. Eliceiri, J. G. Knittel, L. Yan, C. T. Rueden, J. G. White, and P. J. Keely, “Collagen density promotes mammary tumor initiation and progression,” BMC Med. 6, (2008).

20. A. Sugizaki, K. Sato, K. Chiba, K. Saito, M. Kawagishi, Y. Tomabechi, S. B. Mehta, H. Ishii, N. Sakai, M. Shirouzu, T. Tani, and S. Terada, “POLArIS, a versatile probe for molecular orientation, revealed actin filaments associated with microtubule asters in early embryos,” Proc. Natl. Acad. Sci. 118(11), (2021).

21. L. Ravotto, L. Duffet, X. Zhou, B. Weber, and T. Patriarchi, “A Bright and Colorful Future for G-Protein Coupled Receptor Sensors,” Front. Cell. Neurosci. 14, 67 (2020).

22. F. I. Rosell and S. G. Boxer, “Polarized absorption spectra of green fluorescent protein single crystals: Transition dipole moment directions,” Biochemistry 42(1), 177–183 (2003).

23. Y. Chen, X. Song, S. Ye, L. Miao, Y. Zhu, R. G. Zhang, and G. Ji, “Structural insight into enhanced calcium indicator GCaMP3 and GCaMPJ to promote further improvement,” Protein Cell 4(4), 299–309 (2013).

24. R. D. Roorda, T. M. Hohl, R. Toledo-Crow, and G. Miesenböck, “Video-rate nonlinear microscopy of neuronal membrane dynamics with genetically encoded probes,” J. Neurophysiol. 92(1), 609–621 (2004).

25. J. Lazar, A. Bondar, S. Timr, and S. J. Firestein, “Two-photon polarization microscopy reveals protein structure and function,” Nat. Methods 2011 88 8(8), 684–690 (2011).

26. D. J. Dummer, S. G. Kaplan, L. M. Hanssen, A. S. Pine, and Y. Zong, “High-quality Brewster’s angle polarizer for broadband infrared application,” Appl. Opt. 37(7), 1194 (1998).

27. N. Ji, J. C. Magee, and E. Betzig, “High-speed, low-photodamage nonlinear imaging using passive pulse splitters,” Nat. Methods 5(2), 197–202 (2008).

28. J. M. Kralj, A. D. Douglass, D. R. Hochbaum, D. Maclaurin, and A. E. Cohen, “Optical recording of action potentials in mammalian neurons using a microbial rhodopsin.,” Nat. Methods 9(1), 90–5 (2012).

29. W. Bloxham, D. Brinks, S. Kheifets, and A. E. Cohen, “Linearly polarized excitation enhances signals from fluorescent voltage indicators,” bioRxiv 2021.07.21.453006 (2021).

30. K. Sharma, D. K. Fong, and A. M. Craig, “Postsynaptic protein mobility in dendritic spines: long-term regulation by synaptic NMDA receptor activation,” Mol Cell Neurosci 31(4), 702–712 (2006).

